# Genotypic Diversity of Hepatitis B and Delta Viruses in Peru: A Phylogenetic Study Using Next-Generation Sequencing

**DOI:** 10.1101/2025.07.25.666658

**Authors:** Johanna Balbuena-Torres, Ronnie Gavilan-Chavez, Fany Cárdenas-Bustamante, Manuel Terrazas-Aranibar, Maria Martinez-Gago, César Cabezas

## Abstract

**Objective:** To characterize the molecular epidemiology and genomic diversity of hepatitis B virus (HBV) and hepatitis delta virus (HDV) in Peru using next-generation sequencing (NGS).

**Materials and Methods:** A cross-sectional observational study was conducted using 209 serum/plasma samples from various regions of Peru, stored at the National Institute of Health. Detection of HBV and HDV was performed by PCR and RT-PCR, respectively. Nucleic acids were purified and quantified for library preparation and sequencing using the Illumina platform. Sequencing reads were aligned to reference genomes to determine genotypes, subgenotypes, and clinically relevant mutations. Geneious Prime was used for assembly and phylogenetic analysis, and STATA v.14.0 for statistical analysis.

**Results:** HBV genotype F was the most prevalent (96.1%), with subgenotype F1b being predominant, followed by genotypes H, G, and C. Mutations associated with virulence and immune escape were identified, particularly A1762T/G1764A (66.7%) and A1762T/G1764A + G1896A (45.7%). All HDV sequences corresponded to genotype 3 (HDV-3), which was detected in six regions of the country.

**Conclusions:** The findings confirm the predominant circulation of HBV genotype F and HDV genotype 3 in Peru, with genotypic diversity that may influence clinical outcomes, viral load, and response to treatment. Sustained molecular surveillance programs are recommended in endemic areas to guide control, treatment, and prevention strategies.

## INTRODUCTION

Hepatitis B is one of the leading causes of liver cancer worldwide, accounting for 16.4% of global cases and 3.5% in the Americas in 2020 ^(1)^. The World Health Organization (WHO) estimated that in 2022, there were 1.2 million new hepatitis B virus (HBV) infections and approximately 1.1 million deaths from related complications, mainly liver cirrhosis and hepatocellular carcinoma (HCC)^(2)^. Over 254 million people are chronically infected with HBV, but only 13% have been diagnosed and 3% are receiving treatment ^(2)^. In the Americas, around 5 million people were infected in 2022, of whom 2.2 million had access to treatment ^(3)^.

The prevalence of HBV infection varies geographically and is classified as low (<2%), intermediate (2–7%), or high (≥8%)^(4)^. In Peru, the implementation of the national vaccination program since the 1990s has significantly reduced endemicity, although areas with intermediate levels still persist^(4)^.

HBV is a DNA virus from the *Hepadnaviridae* family, with a circular, partially double-stranded 3.2 kb genome organized into four open reading frames (ORFs) that encode the S, C, P, and X proteins. The mutation rate of HBV is high, partly due to the lack of proofreading exonuclease activity in the viral reverse transcriptase^(5,6)^. Mutations can affect diagnostic efficacy, post-vaccination immune response, and the effectiveness of immunoprophylaxis^(7)^.

Ten HBV genotypes (A–J) and over 40 subgenotypes have been identified, with distinct geographic distributions and clinical relevance^(8,9)^. Genotypes are also associated with varying responses to antiviral treatment, clinical progression, and risk of hepatocarcinogenesis ^(10–13)^.

Hepatitis delta virus (HDV) is a defective RNA virus that requires HBV for replication. Chronic HDV infection represents the most severe form of viral hepatitis, with a high rate of progression to cirrhosis, liver failure, and HCC^(14,15)^. Eight HDV genotypes (HDV-1 to HDV-8) with specific geographic distributions have been identified. HDV-3 predominates in South America, especially in Amazonian regions^(16)^. In Peru, previous studies have documented its circulation in Andean and jungle regions, with prevalence rates of up to 39% among HBsAg-positive Amazonian populations ^(17,18,19)^.

Given the clinical importance of HBV and HDV genotypic variants and mutations, the present study aims to characterize the genomic diversity of these viruses in Peru using next-generation sequencing, to provide evidence that can inform improved prevention and control policies for at-risk populations.

## MATERIALS AND METHODS

### Study Design and Population

A cross-sectional observational cohort study was conducted, comprising 209 serum/plasma samples that tested positive for HBV (HBcAg+), of which 27 were also positive for HDV by real-time RT-PCR. The samples were collected between 2021 and 2022 from 20 regions across Peru and are stored in the virotheque of the National Center for Public Health of the National Institute of Health (INS), which serves as the national reference laboratory (see Figure 1).

**Figure 1.**
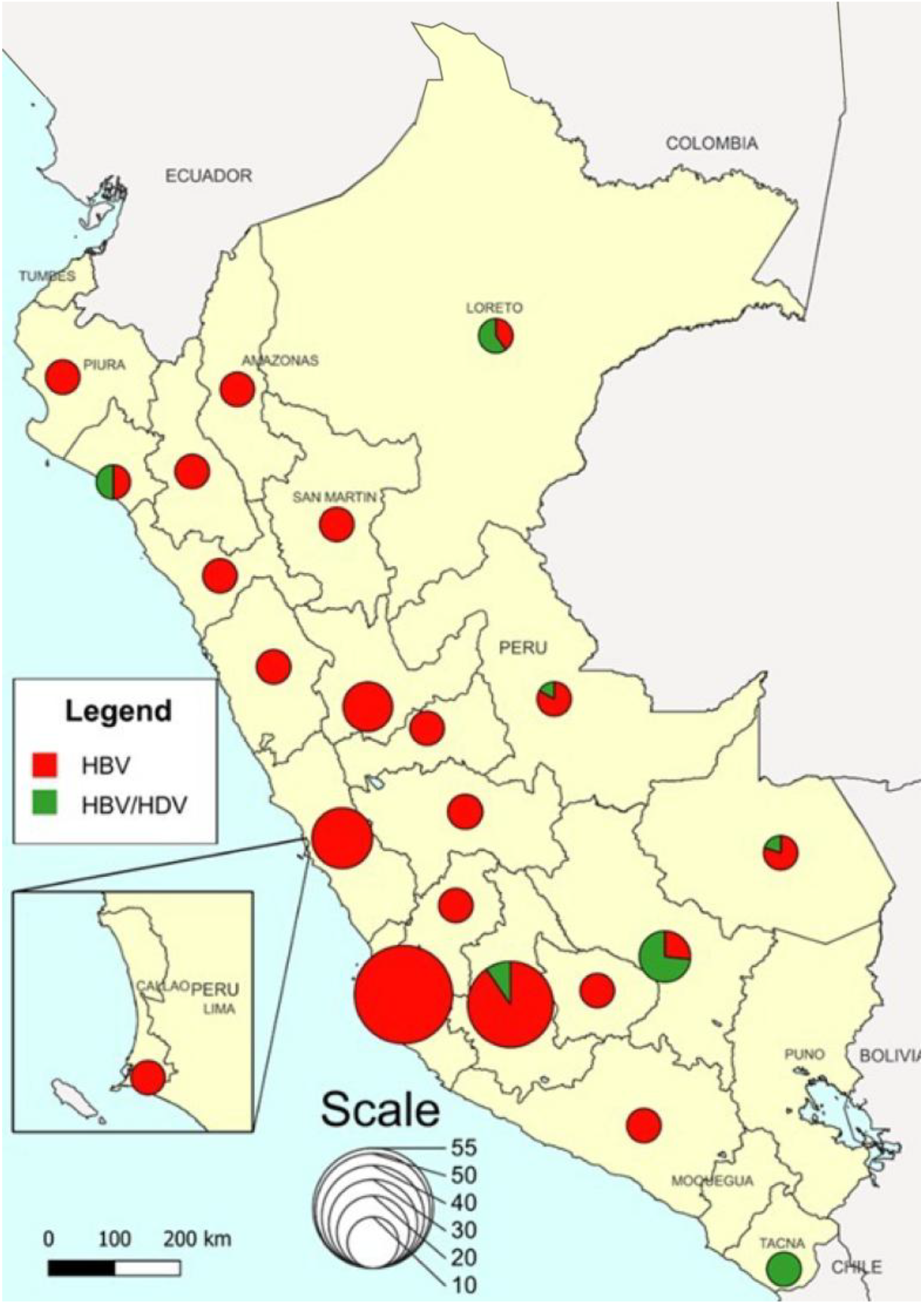
Map of Peru showing the geographic origin of HBV- and HBV/HDV-positive samples included in the study (n = 209). The size of the pie charts indicates the number of from each region, and each segment is color-coded according to the viruses detected. Map created with QGIS Desktop v3.26.3 (http://www.qgis.org).

This study was approved by the Institutional Research Ethics Committee of the INS (code OI-046-21; registration No. 33196-2021). Epidemiological data associated with the samples were analyzed using STATA v.14.0 (StataCorp, College Station, USA).

### Extraction of Viral Genetic Material

A volume of 200 μL from each serum/plasma sample was processed for extraction of HBV viral DNA using the QIAamp DNA Mini Kit (Qiagen, Germany), with a final elution volume of 50 μL. For HDV, 140 μL of viral RNA was extracted using the QIAamp Viral RNA Mini Kit (Qiagen, Germany) and eluted in 60 μL.

### Amplification of the HBV Genome

The complete HBV genome was amplified by nested PCR in two consecutive rounds. The first round used four primer sets (Set 1 to Set 4), and the second round used six pairs (Set 1a to Set 4). Thermal conditions included an initial denaturation at 94°C for 10 minutes, followed by 31 to 35 cycles of denaturation, annealing (50°C to 57°C depending on the primer set), and extension at 72°C for 1 minute. The reaction mix contained primers (2 μL), buffer with Mg^2+^ (5 μL), dNTPs (1 μL), Taq polymerase (1 U), and viral DNA (5 μL in the first round; 3 μL in the second). Amplicons were visualized by 1.5% agarose gel electrophoresis. Band sizes are detailed in Table 1.

**Table 1.**
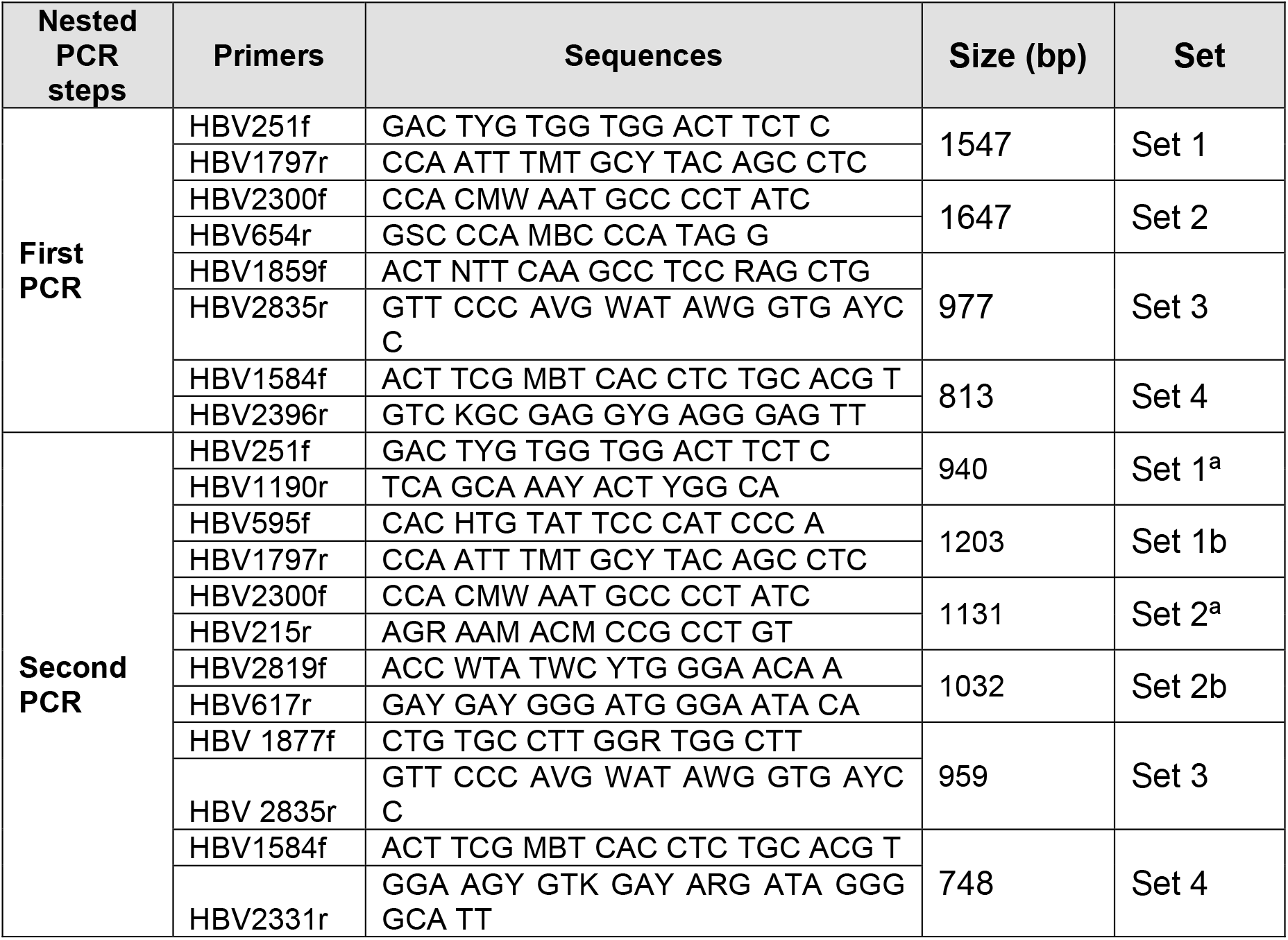
Primer list used for amplification of the hepatitis B virus genome.

### Amplification of the HDV Genome

The complete HDV genome was amplified by multiplex RT-PCR using two primer pools (Pool A and B), each containing two primer pairs. The reaction mix included 5 µL of Capital qPCR Probe Mix 4X, 1 µL of reverse transcriptase with RNase inhibitor (Biotechrabbit), 2 µL of primer pool (5 µM), 7 µL of ultrapure water, and 5 µL of denatured viral RNA (65°C). Thermal cycling involved reverse transcription at 45°C for 60 minutes, followed by touchdown PCR cycles and a final extension at 68°C for 5 minutes. Products were visualized in 1.5% agarose gel. Expected amplicon sizes are listed in Table 2.

**Table 2.**
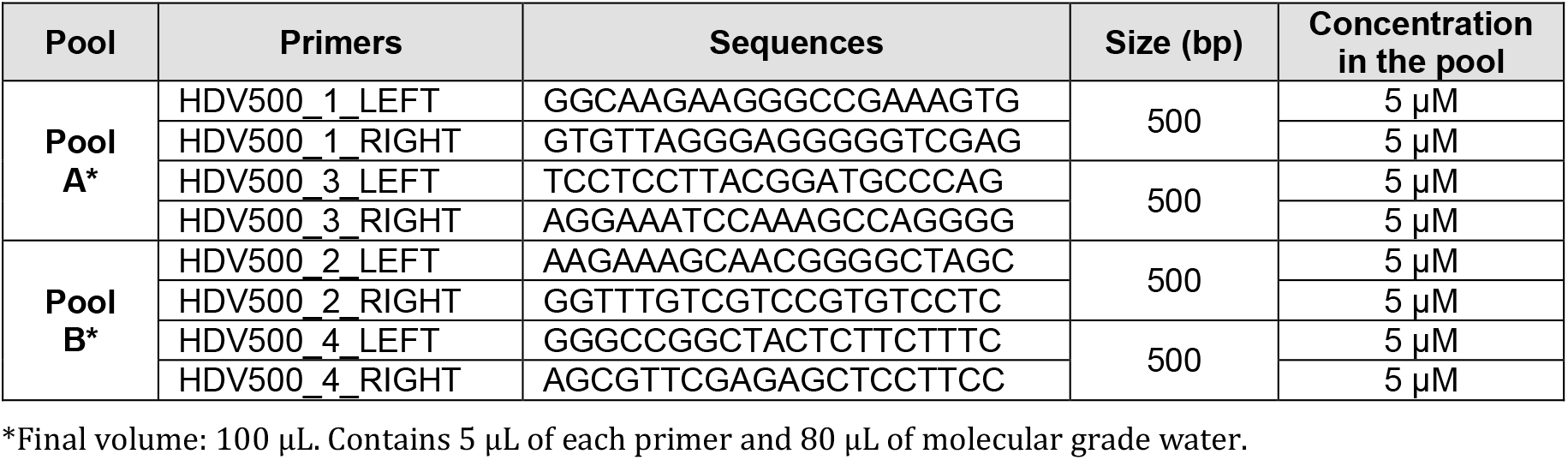
Primer list used for amplification of the hepatitis delta virus genome.

### Library Preparation and Sequencing

Amplicons were purified with AMPure magnetic beads (Beckman Coulter, USA) and quantified using Qubit 3.0 (Invitrogen, Malaysia). Libraries were prepared using the Nextera XT DNA Library Preparation Kit (Illumina, USA) and sequenced on the MiSeq platform (Illumina, USA) with paired-end reads (2 × 300 bp).

### Quality Control and Genome Assembly

The quality of raw reads was assessed using FastQC v0.11.5. Adapters and low-quality bases were trimmed with Trimmomatic v0.38. Genomes were assembled by mapping against HBV reference strain BD14999 (Acc. AY311370) and HDV (Acc. NC_001653.2) using Geneious R11.1 (Dotmatics, UK). Sequencing data were deposited in GenBank under Bioproject PRJNA1253094.

### Phylogenetic Analysis and Genotype Determination

For HBV, sequences with >95% coverage of the S gene were selected, along with 371 reference sequences from GenBank representing all 10 known genotypes. For HDV, sequences with >95% coverage of the full genome were used, along with 86 reference sequences covering the eight known genotypes. Alignments were performed with MAFFT v7, and maximum likelihood phylogenetic trees were constructed using RAxML v8.0 (GTR+G model, 1000 bootstrap replicates). Genotype and subgenotype assignments were validated with HBV-GLUE (for HBV), and results were visualized with iTOL v5.

### Evolutionary Analysis of HBV Subgenotype F1b

BEAST v2.5 was used to estimate the evolutionary rate and demographic history of subgenotype F1b. An exponential coalescent population model, an uncorrelated relaxed molecular clock, and the GTR+G substitution model were applied. MCMC chains were run for 50 million iterations with a 10% burn-in. Convergence was assessed using Tracer v1.7, and a demographic reconstruction with 95% highest posterior density (HPD) intervals was generated.

### Detection of Mutations Associated with Virulence and Antiviral Resistance

HBV sequences were compared with the reference strain BD14999 to identify mutations in the S (immune and diagnostic escape), C and X (virulence), and P (antiviral resistance) genes, based on public databases and current literature^(20)^.

## RESULTS

### Genomic Analysis and Genotypic Diversity of HBV and HDV

Out of the 209 samples processed for HBV genome sequencing, 80 were excluded due to contamination, low quality, or insufficient coverage (<50%). A total of 129 high-quality genomes were obtained, with an average of 366,710 paired-end reads and fragment lengths ranging from 35 to 301 nucleotides. The mean mapping coverage of the genomes was 97.5%. Sequences were obtained from 20 regions of Peru, with the highest frequencies from Ayacucho (n=37), Ica (n=28), and Lima (n=20). The epidemiological characteristics of the patients are shown in Table 3.

**Table 3.** Epidemiological characteristics of HBV-positive patients whose samples were subjected to whole-genome sequencing.

| Region | Sex |  | Age group |  |  | Viral load |  |  |  | Total |
| --- | --- | --- | --- | --- | --- | --- | --- | --- | --- | --- |
|  | Female | Male | 18-29 years | 30-59 years | Older 60 years | Low | Low to Moderate | High | Very High |  |
| Amazonas | 0 | 1 | 0 | 1 | 0 | 1 | 0 | 0 | 0 | 1 |
| Ancash | 2 | 3 | 1 | 4 | 0 | 0 | 3 | 1 | 1 | 5 |
| Apurimac | 3 | 2 | 1 | 4 | 0 | 0 | 5 | 0 | 0 | 5 |
| Arequipa | 0 | 1 | 0 | 0 | 1 | 0 | 0 | 0 | 1 | 1 |
| Ayacucho | 26 | 11 | 14 | 23 | 0 | 2 | 19 | 12 | 4 | 37 |
| Callao | 0 | 1 | 0 | 0 | 1 | 0 | 0 | 0 | 1 | 1 |
| Cusco | 0 | 1 | 1 | 0 | 0 | 0 | 1 | 0 | 0 | 1 |
| Huancavelica | 1 | 0 | 0 | 1 | 0 | 0 | 1 | 0 | 0 | 1 |
| Huánuco | 3 | 0 | 1 | 2 | 0 | 0 | 1 | 2 | 0 | 3 |
| Ica | 18 | 10 | 6 | 18 | 4 | 8 | 11 | 0 | 3 | 28 |
| Junín | 5 | 1 | 1 | 5 | 0 | 0 | 4 | 2 | 0 | 6 |
| La Libertad | 2 | 2 | 1 | 2 | 0 | 0 | 2 | 1 | 1 | 4 |
| Lambayeque | 2 | 0 | 0 | 2 | 0 | 1 | 1 | 0 | 0 | 2 |
| Lima | 9 | 11 | 7 | 11 | 2 | 0 | 10 | 3 | 7 | 20 |
| Loreto | 1 | 1 | 1 | 1 | 0 | 1 | 0 | 1 | 0 | 2 |
| Madre de Dios | 2 | 2 | 2 | 2 | 0 | 3 | 1 | 0 | 0 | 4 |
| Pasco | 2 | 1 | 0 | 3 | 0 | 2 | 0 | 0 | 0 | 3 |
| Piura | 1 | 1 | 0 | 2 | 0 | 0 | 1 | 1 | 0 | 2 |
| San Martín | 1 | 1 | 0 | 2 | 0 | 0 | 2 | 0 | 0 | 2 |
| Ucayali | 1 | 0 | 1 | 0 | 0 | 0 | 1 | 0 | 0 | 1 |
| **Total general** | **79** | **50** | **37** | **83** | **8** | **18** | **63** | **23** | **18** | **129** |

Viral load (Very low: <2,000 IU/mL, Low to moderate: 2,000 – 20,000 IU/mL )
High: >20,000-10<sup>6</sup> IU/mL, Very high: >10<sup>6</sup> IU/mL)

Of the genomes analyzed, four different HBV genotypes were identified. Genotype F was predominant (n=124; 96.1%), followed by genotypes H (n=3; 2.3%), G (n=1; 0.8%), and C (n=1; 0.8%). Within genotype F, subgenotypes F1b (n=115; 89.1%), F3 (n=5; 3.9%), and F4 (n=4; 3.1%) were detected. The only genotype C sample was classified as subgenotype C11. The regional distribution of genotypes is summarized in Table 4.

**Table 4.** Distribution of HBV genotypes by region in Peru.

| Department | HBV genotype |  |  |  | Grand Total |
| --- | --- | --- | --- | --- | --- |
|  | C | F | G | H |  |
| Amazonas | 0 | 1 | 0 | 0 | 1 |
| Ancash | 0 | 5 | 0 | 0 | 5 |
| Apurimac | 0 | 4 | 0 | 1 | 5 |
| Arequipa | 0 | 1 | 0 | 0 | 1 |
| Ayacucho | 0 | 36 | 0 | 1 | 37 |
| Callao | 0 | 1 | 0 | 0 | 1 |
| Cusco | 0 | 1 | 0 | 0 | 1 |
| Huancavelica | 0 | 1 | 0 | 0 | 1 |
| Huánuco | 0 | 3 | 0 | 0 | 3 |
| Ica | 0 | 28 | 0 | 0 | 28 |
| Junín | 0 | 6 | 0 | 0 | 6 |
| La Libertad | 0 | 4 | 0 | 0 | 4 |
| Lambayeque | 0 | 2 | 0 | 0 | 2 |
| Lima | 1 | 17 | 1 | 1 | 20 |
| Loreto | 0 | 2 | 0 | 0 | 2 |
| Madre de Dios | 0 | 4 | 0 | 0 | 4 |
| Pasco | 0 | 3 | 0 | 0 | 3 |
| Piura | 0 | 2 | 0 | 0 | 2 |
| San Martin | 0 | 2 | 0 | 0 | 2 |
| Ucayali | 0 | 1 | 0 | 0 | 1 |
| Grand Total | 1 | 124 | 1 | 3 | 129 |

### HDV Genotypic Characterization

Of the 27 HDV samples sequenced, 9 were excluded due to contamination or low quality. The remaining 18 high-quality sequences came from six regions: Cusco (n=10; 55.6%), Ayacucho (n=2), Loreto (n=2), Tacna (n=2), Madre de Dios (n=1), and Ucayali (n=1). All samples were classified as genotype HDV-3 (100%). In two cases, co-infection with HBV was confirmed, both involving genotype F.

### Phylogenetic Analysis of HBV and HDV

The HBV S gene phylogenetic tree showed a clear geographic clustering. Peruvian genotype F samples, regardless of subgenotype, were closely related to strains from Argentina, Colombia, Chile, and Venezuela. The genotype G sample clustered with Indonesian strains, while genotype C grouped with strains from Europe and North America. The three genotype H samples formed a divergent branch, close to genotype F (Figure 2).

**Figure 2.**
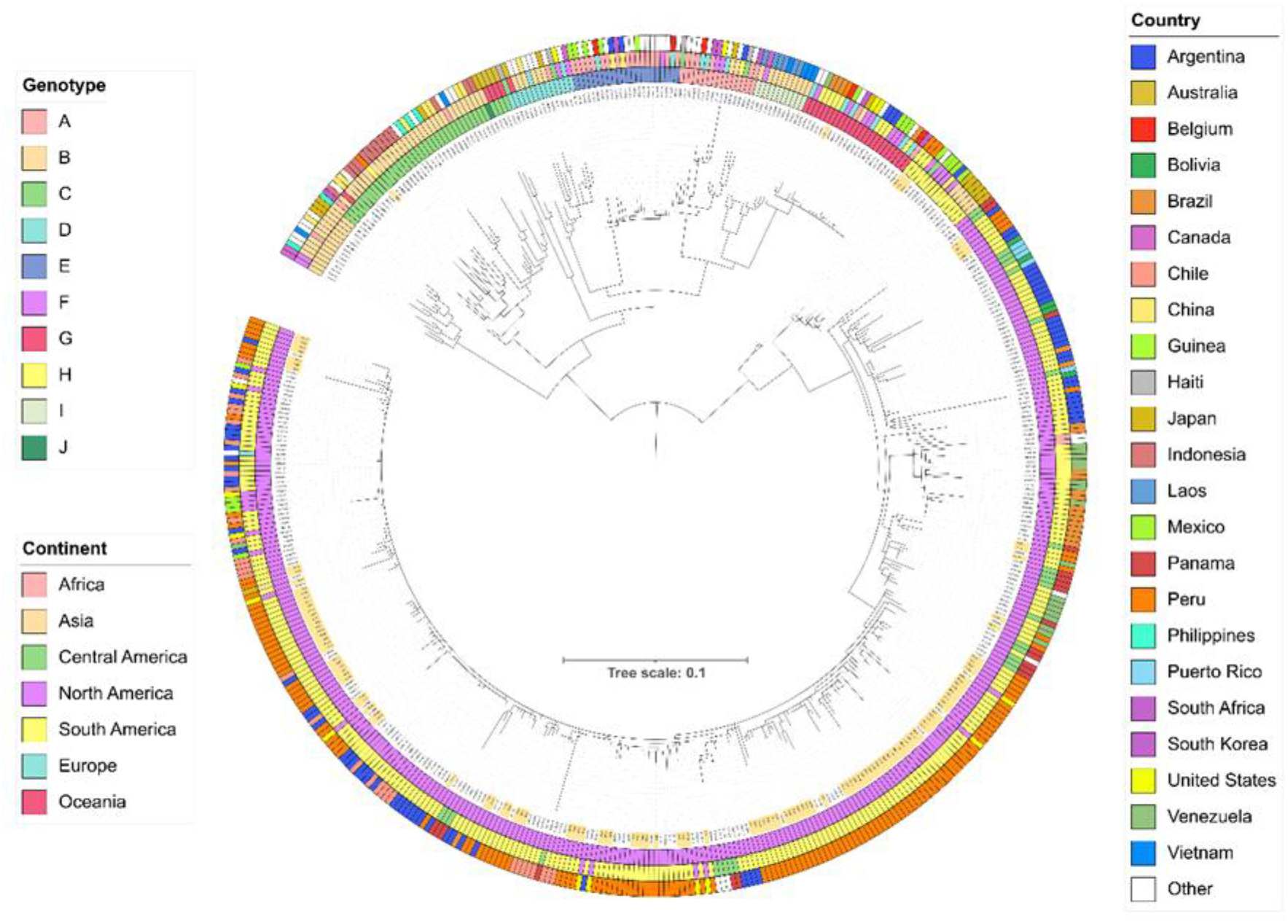
Maximum likelihood phylogenetic tree of the HBV S gene inferred using the GTR+G model with 1000 bootstrap replicates. The inner ring denotes sample genotypes, the middle ring indicates continent of origin, and the outer ring specifies country. The 129 Peruvian samples are marked in orange.

The phylogenetic analysis of complete HDV genomes showed that all Peruvian HDV-3 sequences clustered closely with strains from Brazil and Venezuela, confirming shared regional circulation (Figure 3).

**Figure 3.**
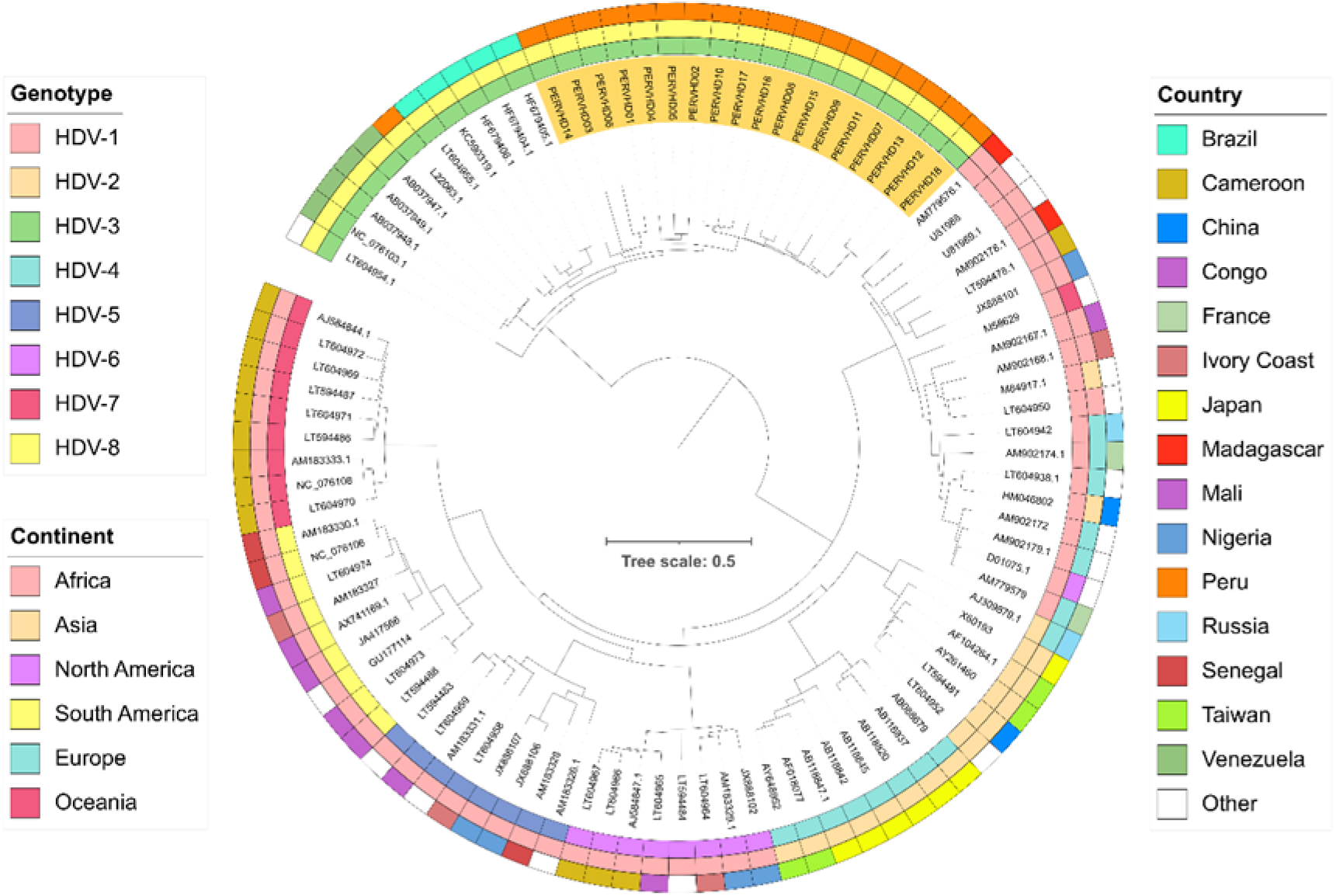
Maximum likelihood phylogenetic tree of the complete HDV genome inferred using the GTR+G model with 1000 bootstrap replicates. The inner ring indicates genotype, the middle ring shows continent of origin, and the outer ring represents countries. The 18 Peruvian samples are marked in orange.

### Evolutionary Analysis of HBV Subgenotype F1b

Bayesian analysis using BEAST v2.5 estimated a nucleotide substitution rate for HBV subgenotype F1b of 1.332 × 10−^4^ substitutions/site/year. The most recent common ancestor (tMRCA) for this subgenotype in South America was dated to around 1960 (95% HPD interval). Demographic reconstruction revealed a sustained population expansion in recent decades (Figure 4).

**Figure 4.**
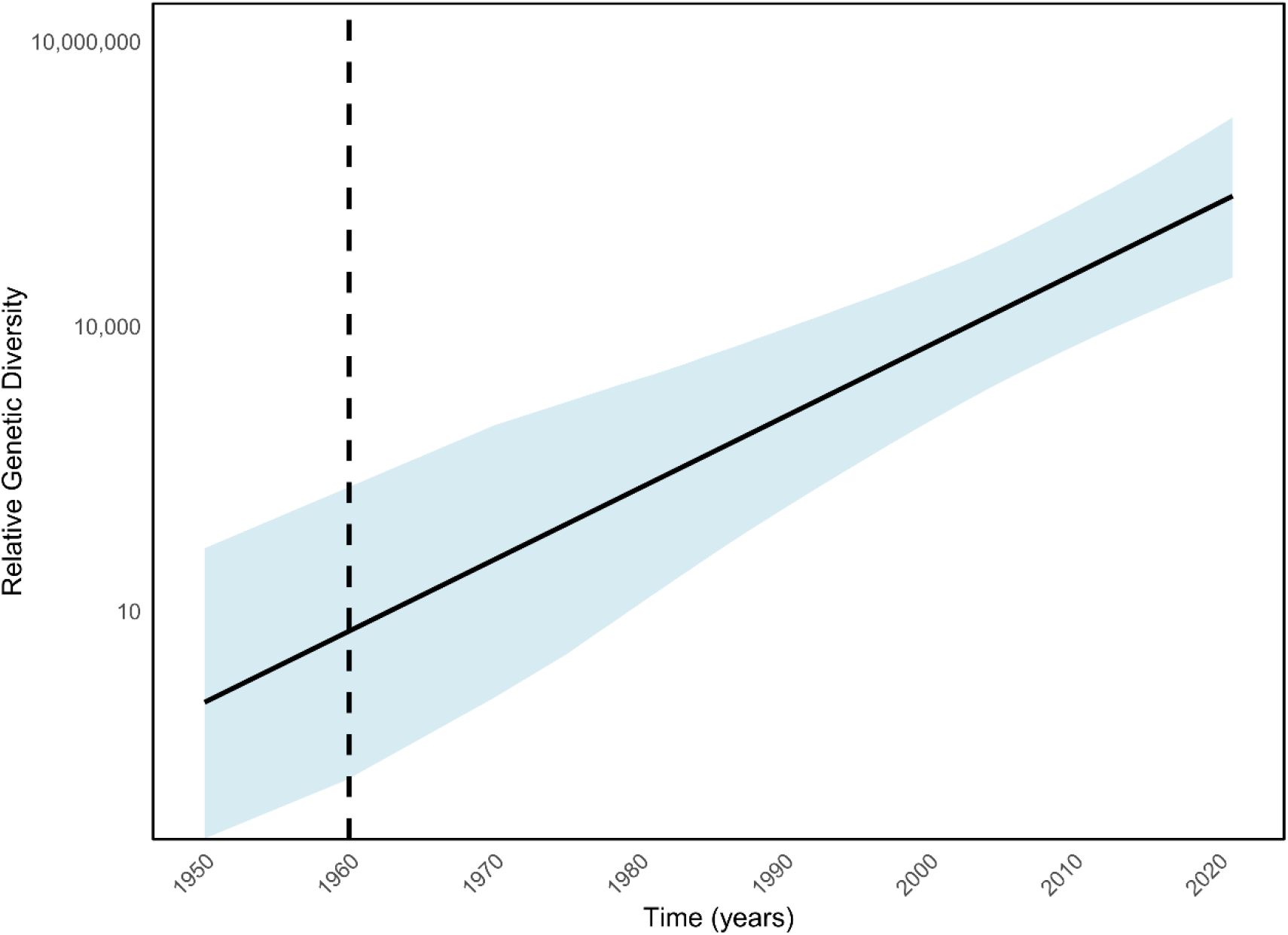
Population dynamics of HBV subgenotype F1b genetic diversity. The Y-axis represents relative genetic diversity, and the X-axis shows chronological time in years. The solid black line indicates the median, with thin lines denoting 95% highest posterior density intervals. Only complete genomes were used in the estimation.

### Mutations Associated with Immune Escape, Diagnostic Failure, and Virulence

Key mutations were detected in the HBV genome. Vaccine escape mutations were identified in six samples (4.7%), most commonly Q129H, Q129R, L109R, and G145R. Three samples (2.3%) had mutations associated with diagnostic escape (P120Q, Q129R, G145R), with Q129R and G145R implicated in both mechanisms.

Multiple mutations associated with severe liver disease progression were also observed. In the basal core promoter (X gene), the most frequent mutations were A1762T (n=93; 72.1%) and G1764A (n=87; 67.4%), followed by T1753V, C1766T, T1768A, and C1653T. In the precore region (pre-C gene), mutations C1858T (n=124; 96.1%), G1896A (n=75; 58.1%), and G1899A (n=13; 10.1%) were found.

Notably, the double mutation A1762T/G1764A (n=86; 66.7%) and its combination with G1896A (n=59; 45.7%) were highly prevalent. These mutations are associated with increased viral replication, HBeAg negativity, and elevated risk of cirrhosis or hepatocellular carcinoma. (Figure 5)

**Figure 5.**
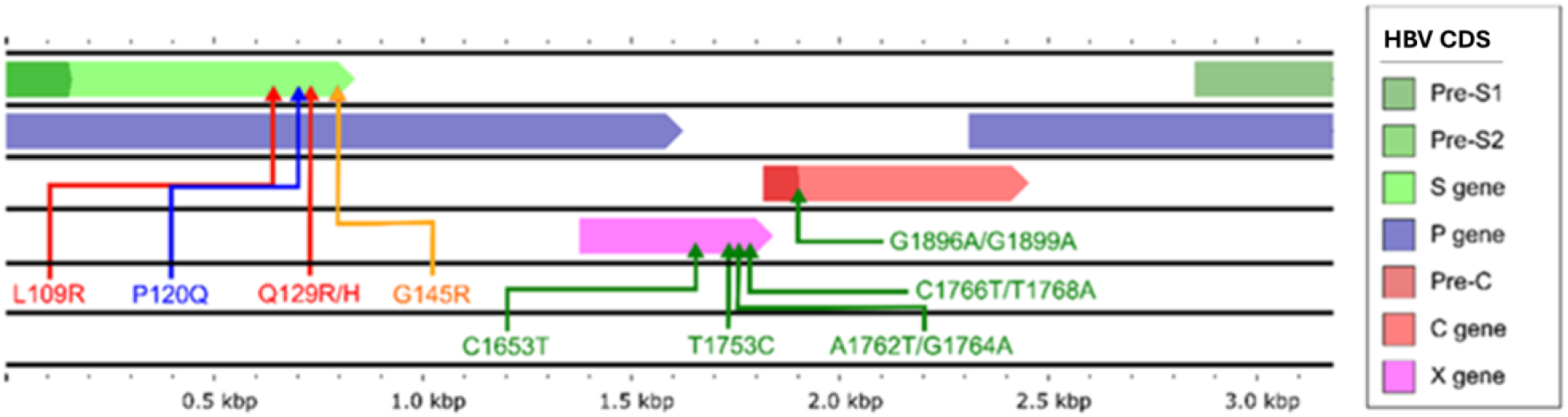
HBV genome structure and major mutations detected in Peruvian samples. Coding regions are shown with colored arrows: precore/core (pre-C, C), polymerase (P), X, and envelope proteins (pre-S1, pre-S2, S). Vaccine escape mutations are in red, diagnostic escape in blue, dual escape in orange, and clinical severity in green.

No mutations associated with antiviral drug resistance were identified, possibly reflecting low selective pressure due to limited use of antiviral therapy in the studied population.

## DISCUSSION

This study provides a comprehensive characterization of the genotypic diversity of hepatitis B virus (HBV) and hepatitis delta virus (HDV) in Peruvian populations using next-generation sequencing (NGS). The findings confirm the predominant circulation of HBV genotype F—particularly subgenotype F1b—and the exclusive presence of HDV genotype 3 among HDV-positive samples.

The high prevalence of genotype F is consistent with previous reports highlighting its predominance in indigenous populations and Amazonian regions of South America. However, the predominance of subgenotype F1b in our study differs in frequency compared to other countries ^(21–23)^. The detection of subgenotypes F3 and F4 in smaller proportions suggests intragenotypic diversity that may have clinical implications, including differences in response to vaccination, antiviral therapy, and disease progression.

All HDV samples were identified as genotype 3 (HDV-3), consistent with prior studies in Peru and neighboring South American countries^(24–25)^. This genotype has been associated with more severe forms of chronic hepatitis and a higher risk of progression to cirrhosis or hepatocellular carcinoma (HCC). Unlike in Brazil where co-infection with HDV-3 and HBV-A has been frequently reported^(25)^ our study found exclusive association with HBV genotype F, revealing epidemiological heterogeneity even among geographically similar settings. The predominance of HDV-3 suggests the possibility that the introduction of HDV into Andean regions may have occurred through human migration and contact with populations from the Amazon, where HBV and HDV endemicity is high^(19)^.

Phylogenetic analysis demonstrated a clear clustering of Peruvian strains with those from other South American countries, supporting the hypothesis of shared regional circulation and potential co-evolution of viral variants. The demographic reconstruction of subgenotype F1b indicated sustained population expansion since the mid-20^th^ century, which may reflect migratory processes or changes in vaccination coverage.

Clinically relevant mutations were identified in key HBV genomic regions. Among them, immune escape mutations (Q129R/H, G145R) and those reducing diagnostic sensitivity (P120Q) were notable. Although their frequency was low (2.3–4.7%), their detection is critical given the scarce genomic data available on Peruvian strains and highlights the need for active surveillance to monitor the emergence and spread of these variants. Other studies have also reported immune escape mutations selected by exposure to nucleos(t)ide analogs^(26)^.

In Peru, tenofovir is currently used to treat chronic HBV infection. This potent antiviral may mitigate the emergence of resistance mutations, though other therapeutic options are more limited. The absence of resistance-associated mutations in this study could be due to the short duration of exposure to antiviral drugs since their introduction in local health services, emphasizing the need for genomic surveillance.

The high frequency of the A1762T/G1764A double mutation (66.7%), which is associated with HBeAg negativity and increased viral replication, is particularly noteworthy. A study from China demonstrated that these mutations are a valuable biomarker for identifying HBsAg-positive males at extremely high risk for hepatocellular carcinoma, with 93.3% of male HCC cases harboring this HBV mutant^(27)^. Likewise, the G1896A mutation (58.1%) in the precore region—also associated with advanced liver disease—appears to have a dual regulatory role in exacerbating HCC severity, as highlighted in another Chinese study^(28)^. These findings underscore the pathogenic potential of circulating HBV variants in Peru.

Chronic hepatitis B (CHB) that is HBeAg-negative is frequently caused by the G1896A mutation in the HBV precore reading frame. Some studies have shown that this stop-codon mutation may enhance the replication of lamivudine-resistant HBV without affecting drug sensitivity in vitro^(29)^. In our study, no mutations associated with resistance to currently used antivirals were found, which may be explained by the limited use of antiviral therapy in the studied population and the resulting low selective pressure. However, spontaneous resistance mutations in untreated patients have been reported in China^(30)^ and Eastern Europe^(31)^.

Among the study’s strengths are the use of NGS for detailed genomic characterization and the wide geographic coverage of the samples. However, limitations include the lack of correlation with clinical data, absence of functional analysis of mutations, and the small number of complete HDV genomes.

In conclusion, HBV genotype F—particularly subgenotype F1b—is the most frequent in Peru, along with less common genotypes H, G, and C. HDV circulates exclusively as genotype 3 (HDV-3), with distribution across both Andean and Amazonian regions. Multiple mutations associated with virulence, HBeAg negativity, and progression to cirrhosis and HCC were identified, notably the high prevalence of the A1762T/G1764A double mutation. Immune and diagnostic escape variants were detected at low frequencies, but their early identification is key for genomic surveillance of HBV in the country. These results highlight the need to implement molecular surveillance programs for HBV and HDV in endemic regions, assess the frequency of escape and resistance mutations, and incorporate these findings into vaccination, early diagnosis, and antiviral treatment policies.

## Funding Source

National Institute of Health (INS), Lima, Peru.

## Conflicts of Interest

The authors declare no conflicts of interest in the publication of this article.

## Notes

### Competing Interest Statement

The authors have declared no competing interest.

## REFERENCES

1. World Health Organization. The Cancers Attributable to Infections [Internet]. 2024 [cited 2025 Jul 9]. Available from: https://gco.iarc.fr/causes/infections

2. World Health Organization. Global Hepatitis Report 2024. Geneva [Internet]. 2024 [cited 2025 Jul 9]. Available from: https://www.who.int/publications/i/item/9789240091672

3. World Health Organization. Global Hepatitis Report 2024: Action for Access in Low- and Middle-Income Countries. Geneva: WHO; 2024. Available from: https://www.who.int/publications/i/item/9789240091672

4. Cabezas C, Trujillo O, Gonzales-Vivanco Á, Benites-Villafane CM, Balbuena-Torres J, Borda-Olivas AO, et al. Seroepidemiology of hepatitis A, B, C, D and E virus infections in the general population of Peru: A cross-sectional study. PLoS One. 2020;15(6):e0234273. doi:10.1371/journal.pone.0234273

5. Assefa A, Getie M, Getie B, Yazie T, Enkobahry A. Molecular epidemiology of hepatitis B virus (HBV) in Ethiopia: A review article. Infect Genet Evol. 2024;122:105618. doi:10.1016/j.meegid.2024.105618

6. Liao F, Xie J, Du R, Gao W, Lan L, Wang M, et al. Replication and expression of the consensus genome of hepatitis B virus genotype C from the Chinese population. Viruses. 2023;15(12):2302. doi:10.3390/v15122302

7. Langat BK, Ochwedo KO, Borlang J, Osiowy C, Mutai A, Okoth F, et al. Genetic diversity, haplotype analysis, and prevalence of hepatitis B virus MHR mutations among isolates from Kenyan blood donors. PLoS One. 2023;18(11):e0289389.

8. Aluora PO, Muturi MW, Gachara G. Seroprevalence and genotypic characterization of HBV among low-risk voluntary blood donors in Nairobi, Kenya. Virol J. 2020;17(1):176. doi:10.1186/s12985-020-01447-2

9. Ward JW, Wanlapakorn N, Poovorawan Y, Shouval D. Hepatitis B vaccines. In: Plotkin S, Orenstein W, Offit P, Edwards KM, editors. Plotkin’s Vaccines. 8th ed. Elsevier; 2023. p. 389–432.e21. doi:10.1016/B978-0-323-79058-1.00027-X

10. Toyé RM, Loureiro CL, Jaspe RC, Zoulim F, Pujol FH, Chemin I. The hepatitis B virus genotypes E to J: the overlooked genotypes. Microorganisms. 2023;11(8):1908. doi:10.3390/microorganisms11081908

11. Kafeero HM, Ndagire D, Ocama P, Kato CD, Wampande E, Walusansa A, et al. Mapping hepatitis B virus genotypes on the African continent from 1997 to 2021: A systematic review with meta-analysis. Sci Rep. 2023;13(1):5723. doi:10.1038/s41598-023-32865-1

12. Arauz-Ruiz P, Norder H, Robertson BH, Magnius LO. Genotype H: a new Amerindian genotype of hepatitis B virus revealed in Central America. J Gen Virol. 2002;83(Pt 8):2059–73.

13. Revill P, Matthews PC, McNaughton AL, McKeating JA, D’Arienzo V, Lumley SF, et al. Insights from deep sequencing of the HB. genome—Unique, tiny, and misunderstood. Gastroenterology. 2019;156(2):384–99.

14. Ilyassova BS, Abzhaparova B, Smailova DS, Bolatov A, Baymakhanov B, Beloussov V, et al. Prevalence and genotypes distribution of hepatitis B and hepatitis delta viruses in chronic liver diseases in Kazakhstan. BMC Infect Dis. 2023;23(1):1030. doi:10.1186/s12879-023-08392-1

15. Colagrossi L, Salpini R, Scutari R, Carioti L, Battisti A, Piermatteo L, et al. HDV can constrain HBV genetic evolution in HBsAg: Implications for the identification of innovative pharmacological targets. Viruses. 2018;10(7):363. doi:10.3390/v10070363

16. Botelho L, Oliveira AL, Villalobos J. Characterization of the genotypic profile of hepatitis delta virus: isolation of HDV genotype-1 in the Western Amazon Region of Brazil. Intervirology. 2015;58(5):166–71. doi:10.1159/000381170

17. Cabezas C, Gotuzzo E, Escamilla J, Irving P. Prevalence of serological markers for viral hepatitis A, B and Delta in apparently healthy schoolchildren in Huanta, Peru. Rev Gastroenterol Peru. 1994;14:123–33.

18. Indacochea S, Gotuzzo E, Delafuente J, Phillips I, Whignal S. High prevalence of hepatitis B and Delta markers in the inter-Andean valley of Abancay. Rev Med Hered. 1991;2(4).

19. Cabezas SC, Suárez JM, Romero CG, Carrillo PC, García MP, Reátegui SJ, et al. Hyperendemicity of hepatitis B and Delta viruses in indigenous peoples of the Peruvian Amazon. Rev Peru Med Exp Salud Publica. 2006;23(2):114–22.

20. Kumar R. Review on hepatitis B virus precore/core promoter mutations and their correlation with genotypes and liver disease severity. World J Hepatol. 2022;14(4):708–18. doi:10.4254/wjh.v14.i4.708

21. von Meltzer M, Vásquez S, Sun J, Wendt UC, May A, Gerlich WH, et al. A new clade of hepatitis B virus subgenotype F1 from Peru with unusual properties. Virus Genes. 2008;37(2):225–30. doi:10.1007/s11262-008-0261-x

22. Moncayo M, Teran E, Gutierrez B, Reyes J, Cortez J, Tobar R, et al. Hepatitis B Virus (HBV) Genotypes in an Ecuadorian Population: A Preliminary Study. Adv Virol. 2024;2024:8823341. doi:10.1155/2024/8823341

23. Santos Alves FAGD, Nogueira Lima FS, Ribeiro JR, Roca TP, Santos AOD, Souza LFB, et al. Genetic diversity of HBV in indigenous populations on the Brazil-Bolivia border. Braz J Infect Dis. 2022;26(5):102700. doi:10.1016/j.bjid.2022.102700

24. Balbuena-Torres J, Santos-Solís L, Navarro-Oviedo R, Cabezas SC. Identification of hepatitis delta virus genotype 3 in Andean and Amazonian communities in Peru. An Fac Med. 2023;84(3):242–8.

25. Crispim MA, Fraiji NA, Campello SC, Schriefer NA, Stefani MM, Kiesslich D. Molecular epidemiology of hepatitis B and delta viruses in the Western Amazon, Brazil. BMC Infect Dis. 2014;14:94. doi:10.1186/1471-2334-14-94

26. Shan M, Shen Z, Sun H, Zheng J, Zhang M. Enrichment of HBV immune-escape mutations during nucleos(t)ide analogue therapy. Antivir Ther. 2017;22(8):717– 20. doi:10.3851/IMP3156

27. Fang ZL, Sabin CA, Dong BQ, Ge LY, Wei SC, Chen QY. HBV A1762T, G1764A mutations as biomarkers for HCC risk in HBsAg-positive males: A prospective study. Am J Gastroenterol. 2008;103:2254–62. doi:10.1111/j.1572-0241.2008.01974.x

28. Zhao B, Qiao H, Zhao Y, Gao Z, Wang W, Cui Y, et al. HBV precore G1896A mutation promotes hepatocellular carcinoma cell growth via ERK/MAPK activation. Virol Sin. 2023;38(5):680–9. doi:10.1016/j.virs.2023.06.004

29. Chen RY, Edwards R, Shaw T, Colledge D, Delaney WE, Isom H, et al. Effect of the G1896A precore mutation on drug sensitivity and replication of lamivudine-resistant HBV in vitro. Hepatology. 2003;37(1):27–35. doi:10.1053/jhep.2003.50012

30. Zhao Y, Wu J, Sun L, Liu G, Li B, Zheng Y, et al. Prevalence of HBV polymerase mutations associated with resistance in treatment-naive CHB patients in Central China. Braz J Infect Dis. 2016;20(2):173–8. doi:10.1016/j.bjid.2015.12.006

31. Salpini R, Svicher V, Cento V, Gori C, Bertoli A, Scopelliti F, et al. Drug-resistance mutations in genotype D HBV-infected patients naïve to antiviral theraPerupy. Antiviral Res. 2011;92(2):382–5. doi:10.1016/j.antiviral.2011.08.013

